## Supplementary material for "Consequences of platelet-educated cancer cells on the expression of inflammatory and metastatic glycoproteins": suppl figures

### Supplementary figures

S1

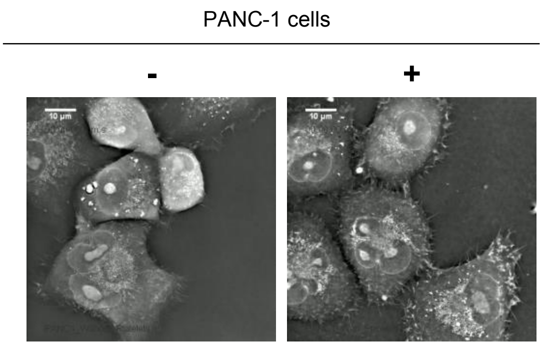


S2

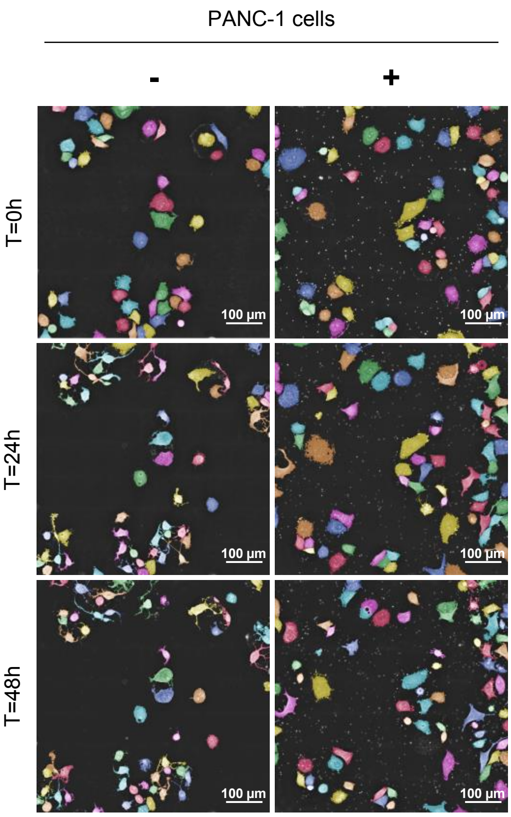


S3

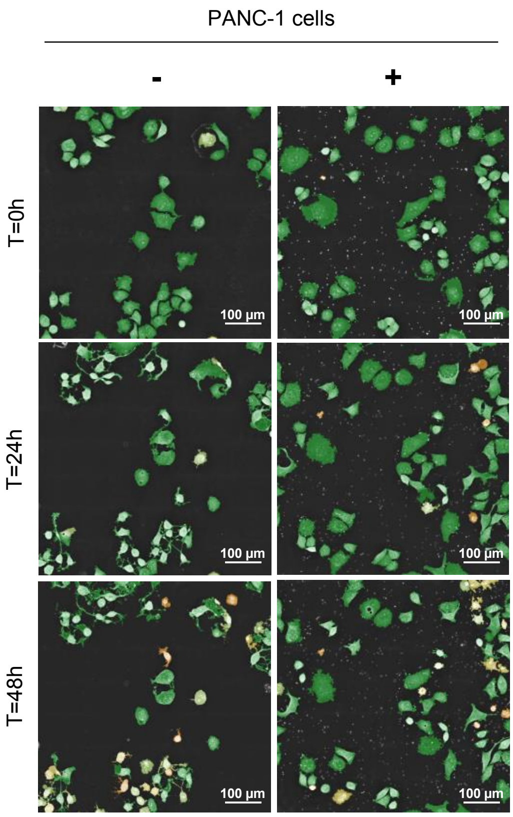


S4

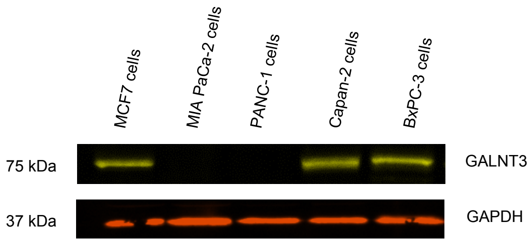


S5

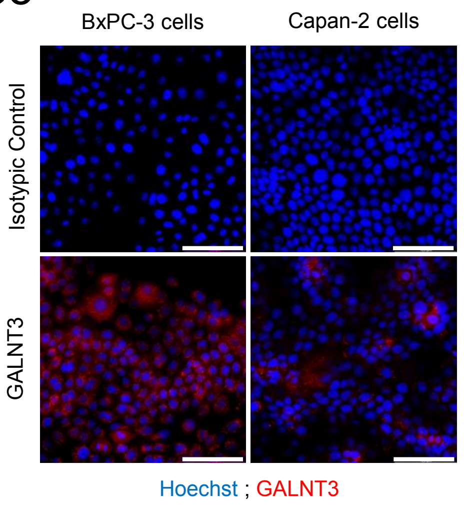


S6

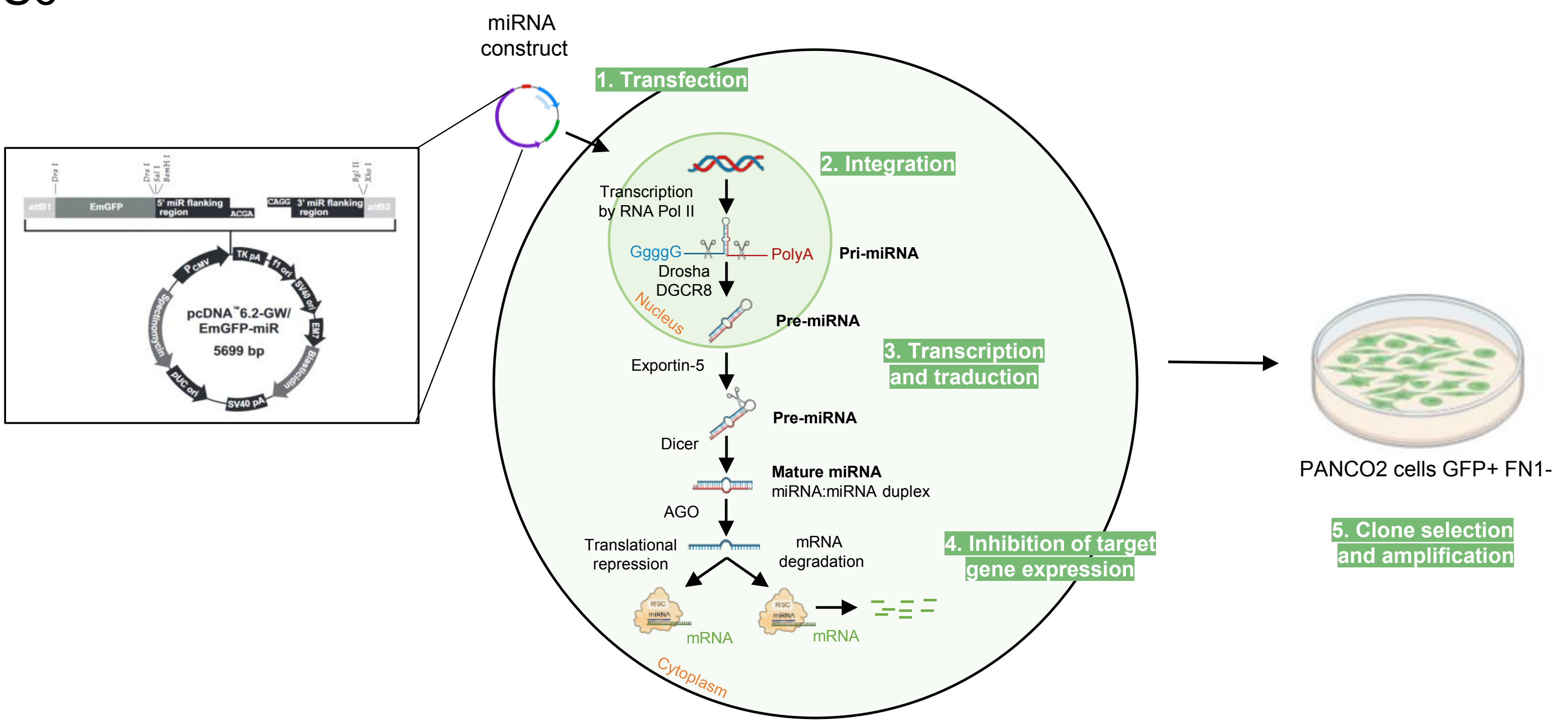

### Supplementary figures

## S7

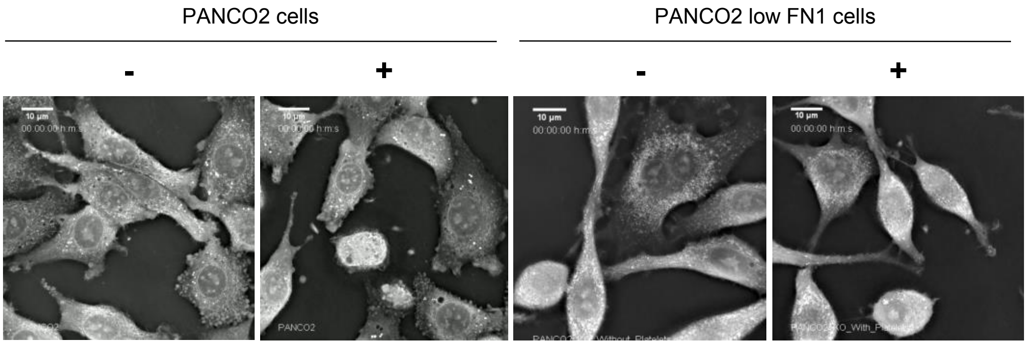

#### Supplementary figures

**(S1)** Representative videos of three independent experiments of PANC-1 cells, in the presence (+) or in absence (-) of platelets over time up to 48 hours (holotomographic microscopy bars, 10 µm). **(S2)** Representative images of three independent experiments of PANC-1 cells in the presence (+) or absence (-) of platelets over time up to 48 hours (holotomographic microscopy bars, 100 µm). Segmentation was performed using "Standard Analysis" module, excluding platelets from the analysis. **(S3)** Representative images of three independent experiments of PANC-1 cells in the presence (+) or absence (-) of platelets over time up to 48 hours (holotomographic microscopy bars, 100 µm). Segmentation was performed using "Cell Death Assay" module, excluding platelets from the analysis. Colors represent live cells (green) or dead cells (yellow). **(S4)** Representative image of GALNT3 (75 kDa) detection by Western Blot in 50 µg of lysate of four different human pancreatic cancer cell lines. MCF7 cell line was used as positive control. Detection of GAPDH was used as a loading control for the experiment. **(S5)** Representative images of indirect GALNT3 (red) detection by immunofluorescence in two human pancreatic cancer cell lines BxPC-3 and Capan-2 using unconjugated anti-human GALNT3 antibody (2 µg/mL) and AF680-conjugated anti-sheep IgG secondary antibody (2 µg/mL) by immunofluorescence in BxPC-3 (bottom, left panel) or Capan-2 cells (bottom, right panel) compared to isotype control (2 µg/mL) revealed with AF680-conjugated anti-sheep IgG secondary antibody in both cases (top, all panels). Cell nuclei (blue) were stained with Hoechst. Images were taken with a 40X objective, 5 images per sample, bars 100 µm. These cell lines were then used as positive controls for GALNT3 expression. **(S6)** Using miRNA transfection, we generated a murine pancreatic cancer line that underexpress fibronectin, called PANCO2 low FN1 cells. **(S7)** Representative videos of three independent experiments of PANCO2 or PANCO2 low FN1 cells during 3 hours of interaction (+) or not (-) with platelets (bars 10 µm).

### Supplementary materials and methods

| Antibody | Supplier | Catalog No. |
|---|---|---|
| **Mouse anti-human CD41, PC7** | Beckman Coulter | #6607115 |
| **Rat anti-mouse GPIb, Dylight 649** | Emfret | #X-649 |
| **Sheep anti-human GALNT3 primary antibody** | Bio-techne | #AF7174 |
| **Sheep IgG Isotype Control** | Thermofisher Scientific | #31243 |
| **Rabbit anti-human FN1 primary antibody** | Cell Signaling Technology | #26836 |
| **Rabbit anti-mouse FN1 primary antibody** | Abcam | #Ab199056 |
| **Rabbit anti-human and mouse GAPDH primary antibody** | Cell Signaling Technology | #2118 |
| **Goat anti-Rabbit IgG secondary antibody, Alexa Fluor Plus 647** | Thermofisher Scientific | #A32733TR |
| **Donkey anti-Sheep IgG secondary antibody, Alexa Fluor 680** | Thermofisher Scientific | #A21102 |
| **Donkey anti-Sheep IgG secondary antibody, Alexa Fluor 488** | Thermofisher Scientific | #A11015 |
| **Hoechst 33342** | Thermofisher Scientific | #H1399 |

**Table 1 :** References of antibodies used

| Lectin | Supplier | Catalog No. |
|---|---|---|
| **VVL/VVA – Biotinylated** | Glycomatrix | #21510196 |
| **ConA lectin – Biotinylated** | Genetex | #GTX01504 |
| **AAL – Biotinylated** | MyBioSource | #MBS656584 |
| **PHA-E – Biotinylated** | Glycomatrix | #30330017 |
| **Streptavidin, Alexa Fluor 647** | Thermofisher Scientific | #S21374 |

**Table 2 :** References of lectins used
